# Efficiency of Anthelmintic Treatment and Its Effect on Microparasite Dynamics in Wild *Mastomys natalensis*

**DOI:** 10.1101/2025.03.01.640952

**Authors:** Marre van de Ven, Bram Vanden Broecke, Alexis Ribas, Herwig Leirs, Christopher Sabuni, Joachim Mariën

## Abstract

Co-infections between helminths and microparasites can influence disease dynamics with potential public health implications. However, establishing clear cause-and-effect relationships from natural populations remains challenging. One approach to address this is through perturbation experiments, where a specific parasite is selectively reduced to directly observe its impact on co-infecting parasites. While such experiments have been conducted in wild rodent populations, they have not yet been explored in Africa, despite the continent’s importance for emerging infectious diseases and zoonoses. In this study, we investigated potential helminth-microparasite interactions in wild *Mastomys natalensis* in Tanzania by using anthelmintics to reduce helminth infections. We first assessed the efficacy of two anthelmintic treatments, ivermectin and pyrantel pamoate, in wild-caught *M. natalensis*, finding that both treatments significantly reduced gastrointestinal nematodes, with pyrantel pamoate being more effective than ivermectin. Additionally, we examined how helminth reduction influenced microparasite prevalence. Our results show that pyrantel pamoate treatment was associated with a decrease in Morogoro virus seropositivity over time, while anthelmintic treatment had no significant effect on *Bartonella* infection probability.

## 1. Introduction

Co-infections are common in both humans and wildlife, and can lead to complex interactions within the host (Petney et al., 1998; Cox, 2001). These interactions can affect host susceptibility, disease severity and parasite transmission rates (Griffiths et al., 2011, Devi et al., 2021). Among co-infections, interactions between helminths and microparasites (viruses, bacteria and protozoa) are of particular interest due to their impact on public health. In human populations, gastrointestinal helminth infections have been linked to increased infection rates and accelerated disease progression of HIV, malaria and tuberculosis (Brown et al. 2006, Mulu et al. 2013, Zenebe et al. 2023), suggesting that helminths may facilitate microparasite infections. Laboratory experiments in rodents indicate that this facilitation is mediated by helminth-induced activation of the Th2 immune response, which suppresses the Th1 response that typically protects against microparasites (Su et al. 2005, Graham 2007, Pedersen et al. 2007, Noland et al., 2008, Su et al. 2014, L et al. 2014).

Most of our current knowledge about co-infections and their interactions comes from observational field studies and controlled laboratory experiments (Lello et al. 2004, Wilson et al. 2007, Graham et al. 2008, Telfer et al. 2010, Salvador et al. 2011, Pol et al. 2017). However, observational studies alone make it difficult to establish cause-and-effect relationships, as species populations are influenced by numerous environmental and host-related factors (Fenton et al. 2014). Laboratory studies, on the other hand, cannot mimic the complexity of natural conditions due to factors such as using inbred mouse strains, administering high infection doses and providing *ad libitum* feeding, limiting the applicability of their findings to wild populations (Viney et al. 2017). Because of these challenges, there is a growing interest in experimental approaches, particularly perturbation experiments, which offer a way to bridge the gap between observational field studies and controlled laboratory experiments (Ezenwa 2016, Fenton et al. 2019). By selectively removing a specific parasite species from a host population using antiparasite treatments, these experiments allow researchers to assess how its absence affects co-infecting parasites (Bender et al. 1984). Comparing treated and untreated hosts helps determine whether the removed parasite facilitated, competed with, or had no effect on other infections, providing clearer insights into causal relationships while still accounting for some of the complexity of natural systems.

Some studies have applied this approach to wild rodent populations, using anthelmintics (i.e. deworming) to investigate helminth-microparasite interactions. Notably, these studies found that reducing helminths resulted in an increase in microparasite prevalence (Knowles et al. 2013, Pedersen et al. 2013, Sweeny et al. 2020), suggesting that helminth infections suppress rather than facilitate microparasite infections. This contrasts with findings from observational and laboratory studies, highlighting the need for more perturbation experiments, which will allow researchers to investigate the direction of causality in natural environments. While such studies have provided valuable insights, their geographic scope remains limited. Despite Africa’s significance in emerging infectious diseases and zoonoses (Jones et al. 2008), no comparable research has yet been conducted there.

To address this, we conducted a perturbation experiment in Morogoro, Tanzania, using the multimammate mouse (*Mastomys natalensis*) as a model to investigate interactions between helminths and microparasites. *Mastomys natalensis* is a common rodent species in sub-Saharan Africa and its population ecology has been extensively studied due to its role as an agricultural pest species and a carrier of zoonotic pathogens (Leirs et al. 2023), highlighting its public health significance. Previous observational studies found positive associations between helminth infections and the presence of microparasites, including Morogoro arenavirus (MORV), a non-pathogenic rodent-borne virus transmitted through direct contact (Hoffmann et al. 2021), and *Bartonella* species, a bacterium spread by bloodsucking arthropod vectors, such as fleas, lice and ticks (Gutiérrez et al. 2015; Vanden Broecke et al. 2021 & 2023). However, the causal relationship between helminth infections and microparasite dynamics, including MORV and *Bartonella* spp., remains unclear.

Building on these findings, we designed a perturbation experiment to investigate potential interactions between helminths, MORV and *Bartonella* in *M. natalensis*. Given the observed positive associations, we hypothesised that helminth infections modulate the immune system by inducing a Th2 skewed response that suppresses Th1-mediated defences, thereby facilitating microparasite infections. We predicted that experimentally reducing helminth infections with anthelmintics would alleviate this suppression and decrease the prevalence of microparasites. To test this hypothesis, we first conducted a controlled laboratory experiment to evaluate the efficacy of two anthelmintic treatments (ivermectin and pyrantel pamoate) in *M. natalensis* using wild-caught individuals. We then monitored microparasites responses in free-living populations of *M. natalensis* using a capture-mark-recapture experiment.

## 2. Material & methods

### 2.1. Study species

*Mastomys natalensis* is a widely distributed rodent in sub-Saharan Africa, commonly found in agricultural fields and human dwellings (Coetzee, 1975; Chidodo et al. 2020). For over 40 years, *M. natalensis* populations have been extensively studied in Morogoro, Tanzania, where both their population ecology and parasite community are well-characterized (Leirs et al. 2023). Gastro-intestinal helminths identified in this population include *Hymenolepis* sp., *Protospirura muricola*, *Syphacia* sp., *Trichuris mastomysi*, *Gongylonema* sp., *Ptergodermatites* sp., *Raillietina* sp., *Inermicapsifer* sp. and the family Trichostrongylidae (Vanden Broecke et al. 2021). Additionally, microparasites such as Morogoro arenavirus (MORV), *Anaplasma*, *Bartonella*, and *Hepatozoon* have been detected (Vanden Broecke et al. 2021 & 2023). MORV is an Old World arenavirus that is non-pathogenic to humans. Transmission is believed to be mainly horizontal (Mariën et al. 2019), occurring through direct contact or environmental exposure to virus particles (Borremans et al. 2015). Available data suggest that MORV infections are characterized by acute viremia, followed by lifelong antibody persistence, and cause minimal or no pathogenic effects (Borremans et al. 2015; Mariën et al 2019; Hoffman et al. 2021). *Bartonella* spp. are facultative intracellular, gram-negative bacteria primarily transmitted by fleas in wild rodent populations and include zoonotic species capable of causing disease in humans (Gutiérrez et al. 2015). Vanden Broecke et al. (2021 & 2023) found that individuals with antibodies against MORV were significantly more likely to be infected with *P. muricola* and that mice infected with *Bartonella* were more likely to be co-infected with helminths. These findings highlight *M. natalensis* as an excellent model for studying co-infection interactions between helminths and microparasites and their potential implications for disease dynamics.

### 2.2. Experimental setup

To evaluate the efficacy of specific anthelmintics (ivermectin and pyrantel pamoate), we first conducted a controlled laboratory experiment with wild-caught *M. natalensis*. The experiment took place at the Sokoine University of Agriculture in Morogoro, Tanzania, in July 2022. Wild rodents were captured in nearby maize and fallow fields using Sherman LFA live traps (Sherman Live Trap Co., Tallahassee, FL). The traps were set in the late afternoon, baited with a mixture of peanut butter and maize flour, and checked the following morning. After capture, rodents were transported to the Institute of Pest Management (PMC) at SUA, where we recorded the sex, weight and reproductive status of each rodent according to Leirs et al. (1994). Rodents were individually housed in standard laboratory cages (28 x 11.5 x 12 cm) with wood shavings as bedding and given food and water *ad libitum*. They were randomly assigned to one of three groups: ivermectin-treated, pyrantel pamoate-treated, and control, with each group containing 20 rodents. One individual from the control group died during the experiment, reducing the sample size to 19. Treatments were administered twice, with a two-week interval between administrations. To minimise the risk of reinfection, cages were cleaned between administrations. After four weeks, the rodents were humanely euthanised with an overdose of isoflurane followed by cervical dislocation. They were then weighed and dissected. The entire gastrointestinal tract was removed and preserved in absolute ethanol for parasite analysis. Additionally, eye lenses were collected and stored in 10% formaldehyde. These lenses were later extracted, cleaned and dried at 100 °C for 2 hours and weighed to the nearest 0.1 mg to estimate their age (Leirs et al. 1990).

Ivermectin and pyrantel pamoate were chosen to reduce helminths burdens, based on their known efficacy in rodents. Ivermectin is commonly used to reduce nematode infections and is known to affect ectoparasitic infestations (Campbell et al., 1983; Wahid et al. 1989; Pedersen et al. 2015), while pyrantel pamoate specifically targets gastrointestinal nematodes without affecting ectoparasitic species relevant to *Bartonella* transmission (Ferrari et al. 2009). Mice were treated orally by gavage with 2 mg/kg ivermectin (Wahid et al. 1989) or 100 mg/kg pyrantel pamoate (Ferrari et al. 2009). A lower dose of ivermectin was used, as ivermectin can have potential neurotoxic effects in certain mouse species at higher doses (Trailovic et al. 2011; Orzechowski et al. 2012). Juveniles (under 16 g) or visible pregnant females were excluded from the treatments.

Following the controlled laboratory experiment, we conducted a field-based capture-mark-recapture (CMR) study to assess the impact of helminth infections on microparasite prevalence in free-living populations. The CMR was set up in Kiroka (Morogoro, Tanzania; coordinates: 6°49’23.0“S 37°47’57.4“E) from August to December 2022. Three 1-hectare grids were selected, each more than 200 meters apart with natural and artificial barriers between them to minimise movements of individuals between the grids. Within each grid, 100 Sherman LFA live traps were arranged in rows of ten, each spaced five metres apart. Trapping sessions were conducted every two weeks, each lasting for three consecutive nights. Traps were baited as described above, set at dusk, and checked the following morning. Captured rodents were processed individually and data on sex, body weight and reproductive status were noted. Blood samples were collected from the retro-orbital sinus and were stored on Whatmann filter paper for microparasite screening. Newly captured individuals were marked by clipping terminal phalanges to ensure accurate individual identification over time (Leirs et al. 2023). Each rodent was then randomly assigned to receive ivermectin, pyrantel pamoate or no treatment (control group) following the same treatment protocol used in the laboratory experiment. Treatment was repeated upon recapture in the subsequent trapping session. After processing, all rodents were released at their exact capture location. Rodents recaptured within the same trapping session were immediately released without further handling.

### 2.3. Parasitological examination

Helminths were isolated from the gastrointestinal tract according to established protocols (Ribas et al. 2011; Vanden Broecke et al. 2021). Gastrointestinal contents were examined under a stereomicroscope and helminths were identified based on morphological characteristics using a light microscope.

In addition to helminth analysis, blood samples were screened for Morogoro virus (MORV) and *Bartonella* infection to investigate possible interactions between microparasites and helminth presence in the study population. Morogoro virus-specific IgG antibodies (MORVab) were detected using an immunofluorescence assay (Borremans et al. 2014; Mariën et al. 2017). Positive controls were added to each slide and all slides were checked twice. In case of uncertainty, samples were retested. To detect *Bartonella* infection, DNA was extracted from the blood samples and analysed using real-time quantitative PCR (qPCR) targeting the 16S-23S intergenic transcribed spacer (ITS) region. Positive samples were amplified with conventional PCR and the products were sequenced with the forward primer (Dahmana et al. 2020; Vanden Broecke et al. 2023). Sequences were processed in Geneious Prime v2024.0.3, where primers were removed and low-quality regions were trimmed. They sequences were then aligned using Clustal-W within Geneious Prime, incorporating previously identified *Bartonella* isolates from rodents and known *Bartonella* species. A Neighbor-Joining tree was generated using the Tamura-Nei model and visualised with iTOL v7 for phylogenetic analysis. *Bartonella* subgroups were defined based on a 93.9% similarity threshold for the ITS gene, as established by La Scola et al. (2003).

### 2.4. Statistical analyses

All statistical analyses were performed using R (v4.4.1; R Core Team 2024) with the ‘lme4’ package (Bates et al. 2015). To evaluate the effect of treatment on the presence or absence of helminths in mice from the laboratory experiment, we ran generalized linear models (GLMs) with a binomial error distribution. For the main analysis, helminths from the same phylum (nematodes or cestodes) were pooled, but the nematode species with the highest prevalence, *Trichuris mastomysi*, was also analysed separately. In all models, we included the same fixed effects, namely: treatment (control, ivermectin, pyrantel pamoate), sex (female, male) and age in days, along with potential interaction terms between treatment and the other variables. Age was estimated using the equation published in Leirs et al. (1990). For five individuals, eye lens weight measurements were missing. Since body weight is correlated with age, we modeled the relationship between eye lens weight and body weight for individuals with both measurements available and used this model to estimate missing eye lens weights. All animals were under 1 year old (<250 days). Models were simplified using backward stepwise elimination of non-significant terms (p>0.05) beginning with interaction terms, to obtain the minimum adequate model. To assess whether treatment affected the health and well-being of the mice, we analysed changes in body weight using a two-way ANOVA, with treatment and time (before and after treatment) as predictors.

For the capture-mark-recapture study, we analysed the effect of treatment on Morogoro virus-specific antibodies (MORVab) and *Bartonella* infection using generalized linear mixed models (GLMMs). A total of 381 individual *M. natalensis* were captured, out of which 75 were selected for analysis (22 with ivermectin, 22 with pyrantel pamoate and 31 as control). Individuals were included if they had been captured at least three times across different trapping sessions. The first capture was used as time 0 and subsequent time points were calculated relative to this first capture in weeks. This resulted in capture frequencies ranging from 3 to 10 times per individual over an 18-week period (**Supplementary Figure S1**). The highest capture rate occurred within the first four weeks, followed by a gradual decline in the following weeks. To ensure an adequate and balanced sample size, we focused the model analysis on data from the first four weeks. The presence or absence of MORVab and *Bartonella* infection served as response variables and were modelled using a binomial distribution. Fixed effects included treatment, trapping grid (KIR3, KIR4, KIR5), sex, body weight (as a rough proxy for age: Leirs et al. 1990), time (0, 2 or 4 weeks) and the interaction between time and treatment. Missing infection status values at a given time point were imputed based on the last observed status. To account for repeated measurements within individuals, we included individual ID as a random effect.

## 3. Results

### 3.1. Laboratory experiment

Seven helminths species (five nematodes and two cestodes) were identified in the gastrointestinal tract of *M. natalensis* (**Table 1**). Among the nematodes, four genera were identified: *Trichuris mastomysi*, *Syphacia* sp., *Gongylonema* sp. and *Protospirura* sp., along with one family, Trichostrongylidae, for which no specific genera could be distinguished. Among the cestodes, one genus, *Hymenolepis* spp. was identified, as well as one family, Davaineidae. In addition, cysts of the cestode *Taenia* sp. were found in the body cavity.

**Table 1:**
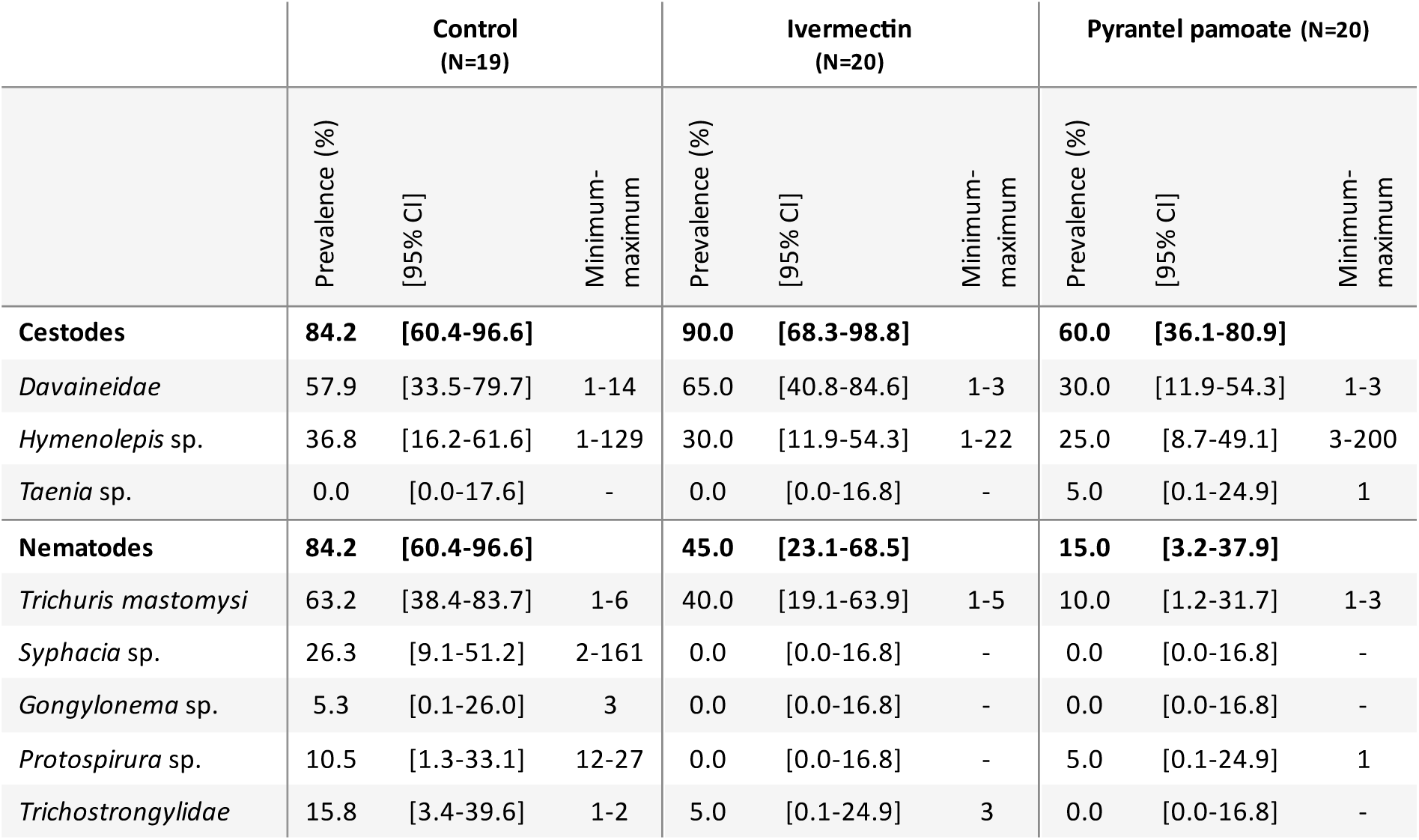
Prevalence, 95% binomial confidence intervals and range (minimum-maximum) of parasite load for each helminth species found in *M. natalensis* during the laboratory experiment following treatment.

|  | Control<br>(N=19) |  |  | Ivermectin<br>(N=20) |  |  | Pyrantel pamoate (N=20) |  |  |
| --- | --- | --- | --- | --- | --- | --- | --- | --- | --- |
|  | Prevalence (%) | [95% CI] | Minimum-maximum | Prevalence (%) | [95% CI] | Minimum-maximum | Prevalence (%) | [95% CI] | Minimum-maximum |
| **Cestodes** | **84.2** | **[60.4-96.6]** |  | **90.0** | **[68.3-98.8]** |  | **60.0** | **[36.1-80.9]** |  |
| *Davaineidae* | 57.9 | [33.5-79.7] | 1-14 | 65.0 | [40.8-84.6] | 1-3 | 30.0 | [11.9-54.3] | 1-3 |
| *Hymenolepis* sp. | 36.8 | [16.2-61.6] | 1-129 | 30.0 | [11.9-54.3] | 1-22 | 25.0 | [8.7-49.1] | 3-200 |
| *Taenia* sp. | 0.0 | [0.0-17.6] | - | 0.0 | [0.0-16.8] | - | 5.0 | [0.1-24.9] | 1 |
| **Nematodes** | **84.2** | **[60.4-96.6]** |  | **45.0** | **[23.1-68.5]** |  | **15.0** | **[3.2-37.9]** |  |
| *Trichuris mastomysi* | 63.2 | [38.4-83.7] | 1-6 | 40.0 | [19.1-63.9] | 1-5 | 10.0 | [1.2-31.7] | 1-3 |
| *Syphacia* sp. | 26.3 | [9.1-51.2] | 2-161 | 0.0 | [0.0-16.8] | - | 0.0 | [0.0-16.8] | - |
| *Gongylonema* sp. | 5.3 | [0.1-26.0] | 3 | 0.0 | [0.0-16.8] | - | 0.0 | [0.0-16.8] | - |
| *Protospirura* sp. | 10.5 | [1.3-33.1] | 12-27 | 0.0 | [0.0-16.8] | - | 5.0 | [0.1-24.9] | 1 |
| *Trichostrongylidae* | 15.8 | [3.4-39.6] | 1-2 | 5.0 | [0.1-24.9] | 3 | 0.0 | [0.0-16.8] | - |

Sex and age had no significant effect on the presence of nematodes or cestodes, nor did treatment effects differ based on sex or age (*p>0.189;* **Supplementary Tables S1 & S3**). Both ivermectin and pyrantel pamoate treatments significantly reduced the prevalence of intestinal nematodes compared to the untreated control group. However, pyrantel pamoate (β=−3.409, s.e.=0.888, *p*<0.001; **Figure 1A**) was more effective than ivermectin (β=−1.875, s.e.=0.773, *p*=0.015) in reducing overall nematode prevalence. This difference in efficacy was also reflected in *T. mastomysi* prevalence, with pyrantel pamoate significantly reducing *T. mastomysi* prevalence (β=−2.736, s.e.=0.884*, p=0.002;* **Figure 1B; Supplementary Table S2**), while ivermectin did not significantly reduce *T. mastomysi* prevalence (β=−0.945, s.e.=0.659*, p=0.152*). No significant treatment effects were observed for cestodes: ivermectin had no significant effect (β=0.523, s.e.=0.975, *p*=0.592) and although pyrantel pamoate showed a reduction, it was not statistically significant (β=−1.269, s.e.=0.777, *p*=0.103). Finally, no significant changes in body weight were observed between the beginning and end of the experiment following treatment (F=0.495, df=2, *p*=0.610).

**Figure 1:**
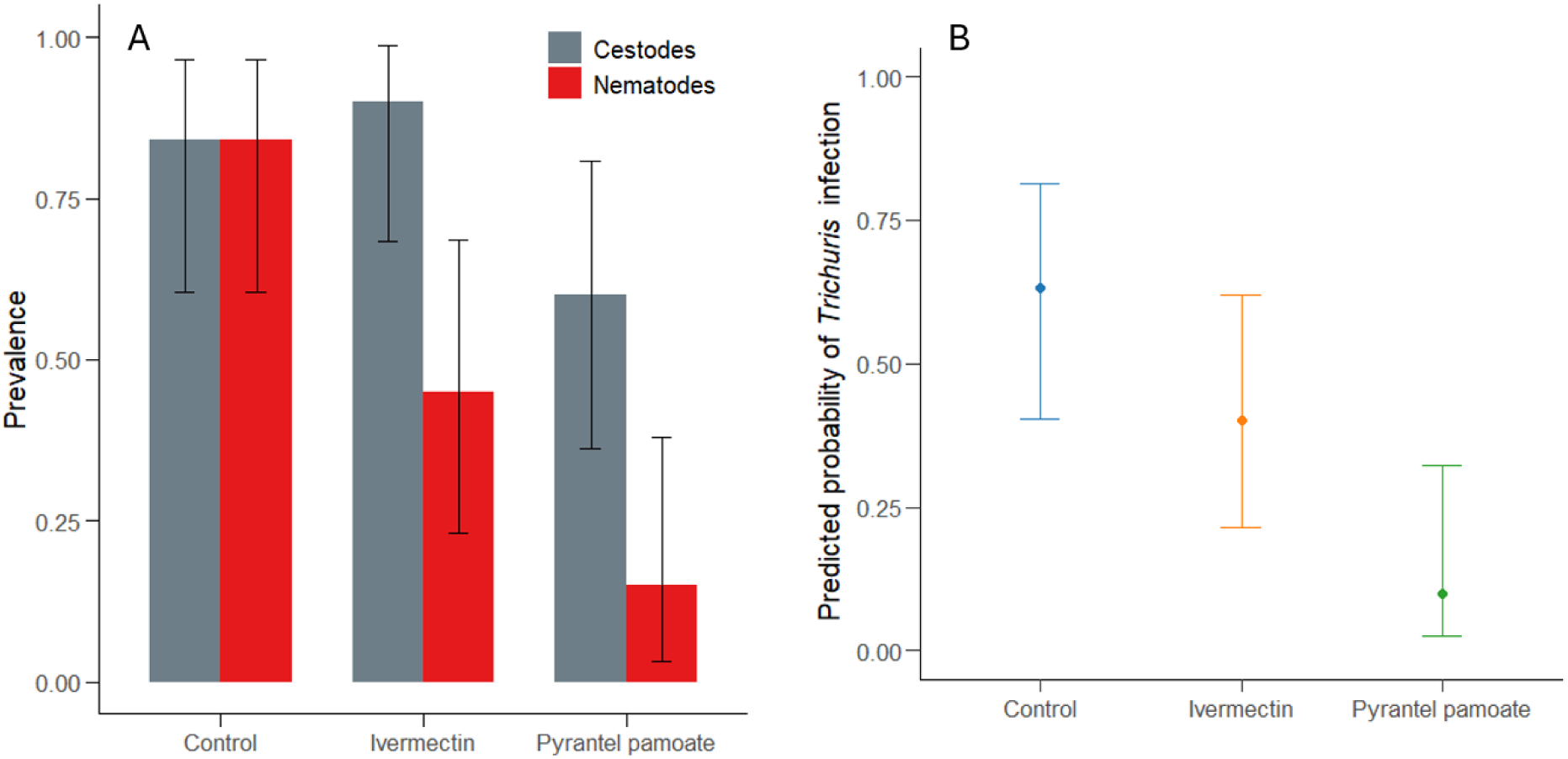
(A) Prevalence of nematodes and cestodes, and (B) predicted probability of *Trichuris mastomysi* infection in *Mastomys natalensis* across treatment groups (control, ivermectin and pyrantel pamoate), with 95% confidence intervals.

### 3.2. Capture-mark-recapture study

#### 3.2.1. Morogoro virus infection dynamics

During the study, 35 of 75 individuals (47%) tested positive for MORVab at least once, and seven individuals were seropositive for all their samples (**Supplementary Figure S2**). In the control group, the probability of seropositivity increased over time, although this trend was not statistically significant (β=0.408, s.e.=0.240, *p*=0.088; **Figure 2A**). In contrast, the pyrantel pamoate group showed a significant negative interaction with time, suggesting that treatment with pyrantel pamoate reduced the likelihood of seropositivity over time compared to the control group (β=−0.829, s.e.=0.384, *p*=0.031). The ivermectin-treated group showed no significant effect on seropositivity over time (β=−0.062, s.e.=0.355, *p*=0.862). Sex and trapping grid had no significant effect either (*p*>0.460: **Supplementary Table S4**), while body weight was a significant predictor, with heavier animals being more likely to test positive for MORVab (β=0.112, s.e.=0.051, *p*=0.028).

**Figure 2:**
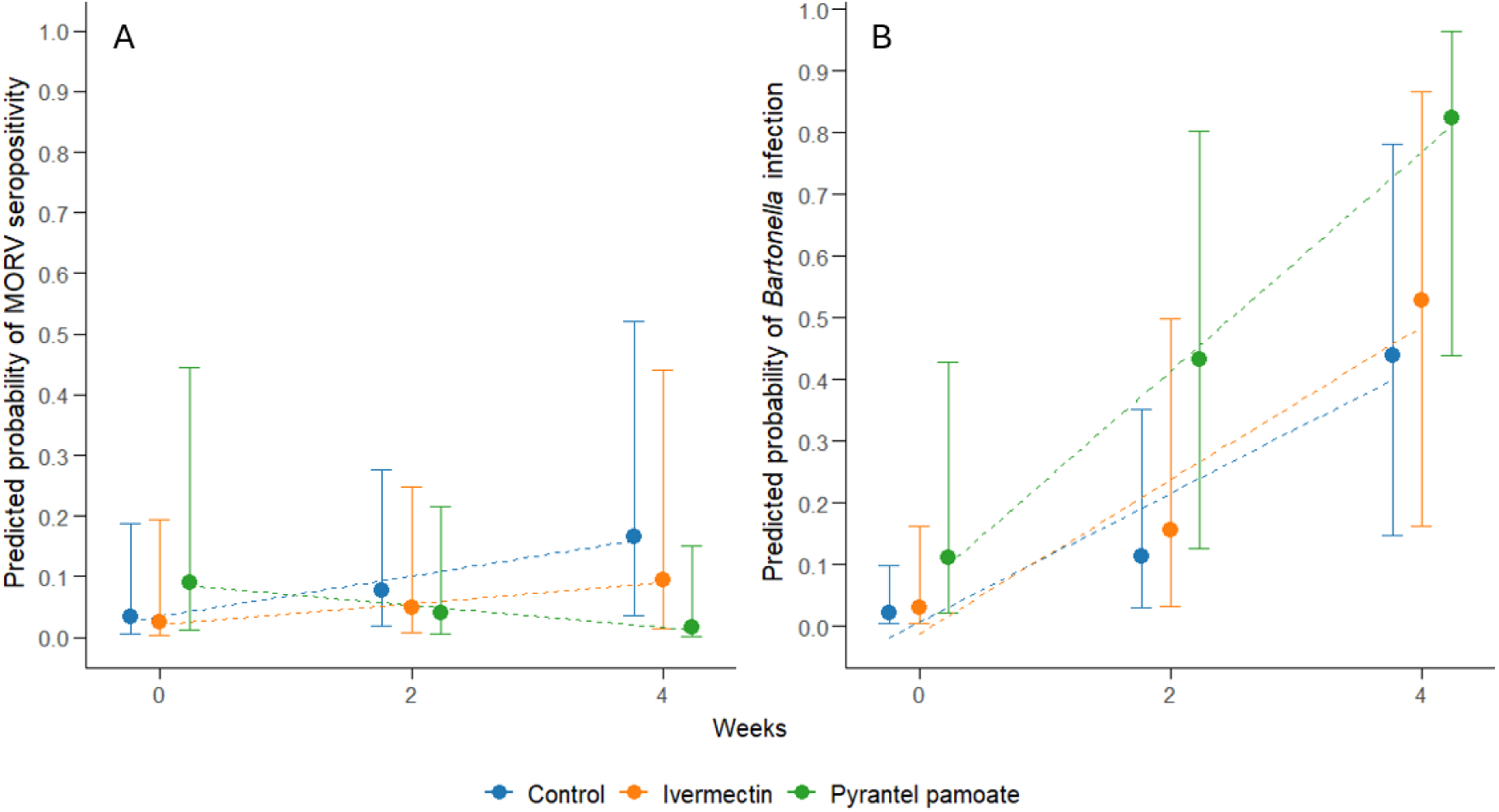
**(A)** Predicted probability of Morogoro virus (MORV) seropositivity and **(B)** *Bartonella* infection over time (weeks) for each treatment group (control, ivermectin and pyrantel pamoate) in a wild *Mastomys natalensis* population. Error bars represent 95% confidence intervals and dashed lines indicate linear trends.

#### 3.2.2. *Bartonella* species diversity

Of the 294 samples screened (each sample representing a biological specimen collected from an individual at a specific time point), 135 (45.9%) tested positive for *Bartonella* based on sequence analysis. We excluded 17 sequences from the phylogenetic analysis due to low quality and were classified as ‘Unspecified’. The remaining sequences clustered into four genogroups and seven subgroups based on sequence identity (**Figure 3**). Genogroups A and B clustered with *Bartonella* sequences previously identified in *Arvicanthis niloticus* from Uganda (GenBank accession no. JX428757 & JX428753; Billeter et al. 2014). Notably, genogroup A was almost exclusively found in KIR5. Genogroup C showed some similarity to *B. tribocorum* (>85% identity, >50% query coverage) but did not cluster with previously identified *Bartonella* isolates. Genogroup D, which accounted for 72.6% of the sequences, showed similarity to *B. elizabethae* (>85 identity, >90% coverage) and shares high identity with sequences previously identified from *Mastomys natalensis* in Tanzania (GenBank accession no. OL984911), *Mastomys erythroleucus* in Senegal (GenBank accession no. MN158196) and other rodents from Uganda, Myanmar and China (Billeter et al. 2014; Dahmana et al. 2020). Subgroup D1 was the most prevalent across all groups (**Supplementary Table S5**).

**Figure 3:**
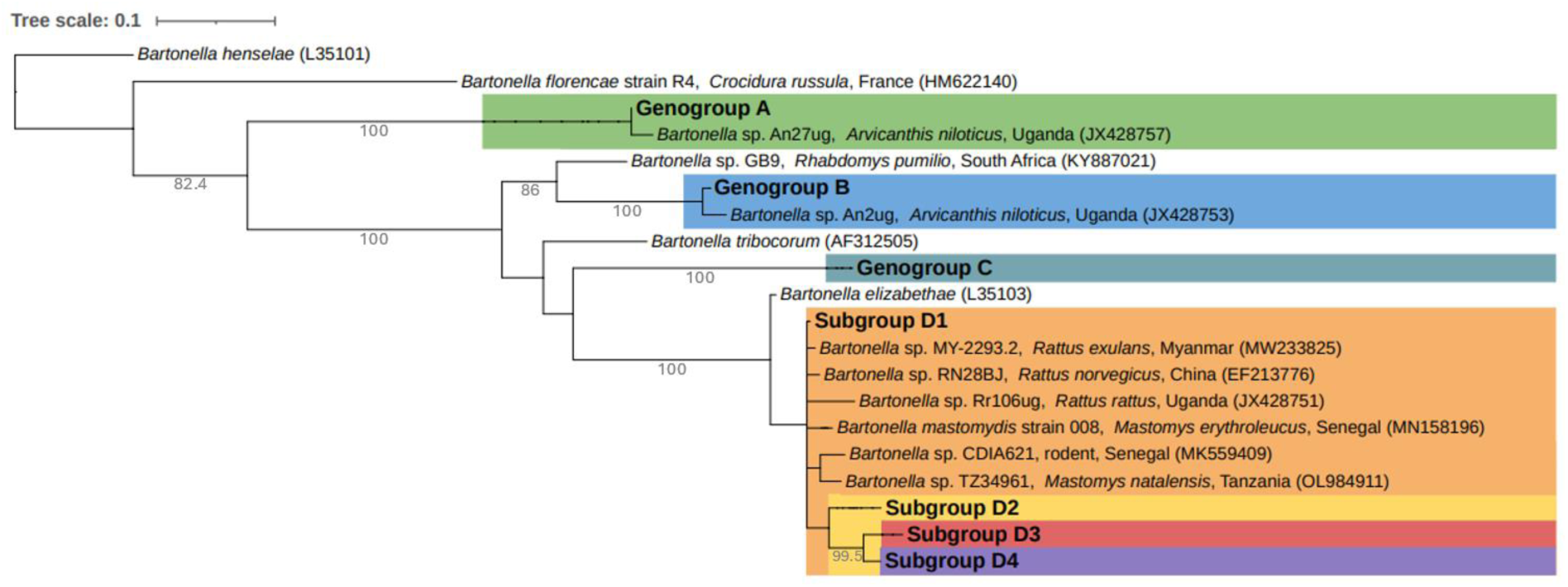
Phylogenetic relationship of *Bartonella* sequences identified in this study. A Neighbor-Joining tree was constructed using the Tamura-Nei model based on ITS gene sequences, with reference sequences from previously identified *Bartonella* isolates in rodents and recognized *Bartonella* species. Bootstrap support values above 70% are shown (1,000 replicates). Subgroups were defined based on a 93.9% similarity cut-off (La Scola et al., 2003). The scale bar represents 0.1 nucleotide substitutions per site.

#### 3.2.3. *Bartonella* infection dynamics

Over the course of the study, 49 out of 75 individuals (65.3%) tested positive for *Bartonella* at least once, with 42 individuals testing positive on multiple occasions (**Figure 4**). Infection with the same subgroup lasted up to eight weeks and some individuals carried different subgroups over time. The probability of *Bartonella* infection increased significantly over time (β=1.065, s.e.=0.281, *p*<0.001). Neither ivermectin nor pyrantel pamoate treatments significantly affected this trend (*p*>0.193). Sex had no significant effect on infection probability (β=−0.118, s.e.=0.787, *p*=0.880; **Supplementary Table S6**). In contrast, body weight and trapping grid were significant predictors: heavier animals were more likely to be infected (β=0.225, s.e.=0.063, *p*=0.003) and individuals from grid KIR4 had a significantly lower probability of infection compared to those from KIR3 (β=−4.234, s.e.=1.430, *p*=0.003).

**Figure 4:**
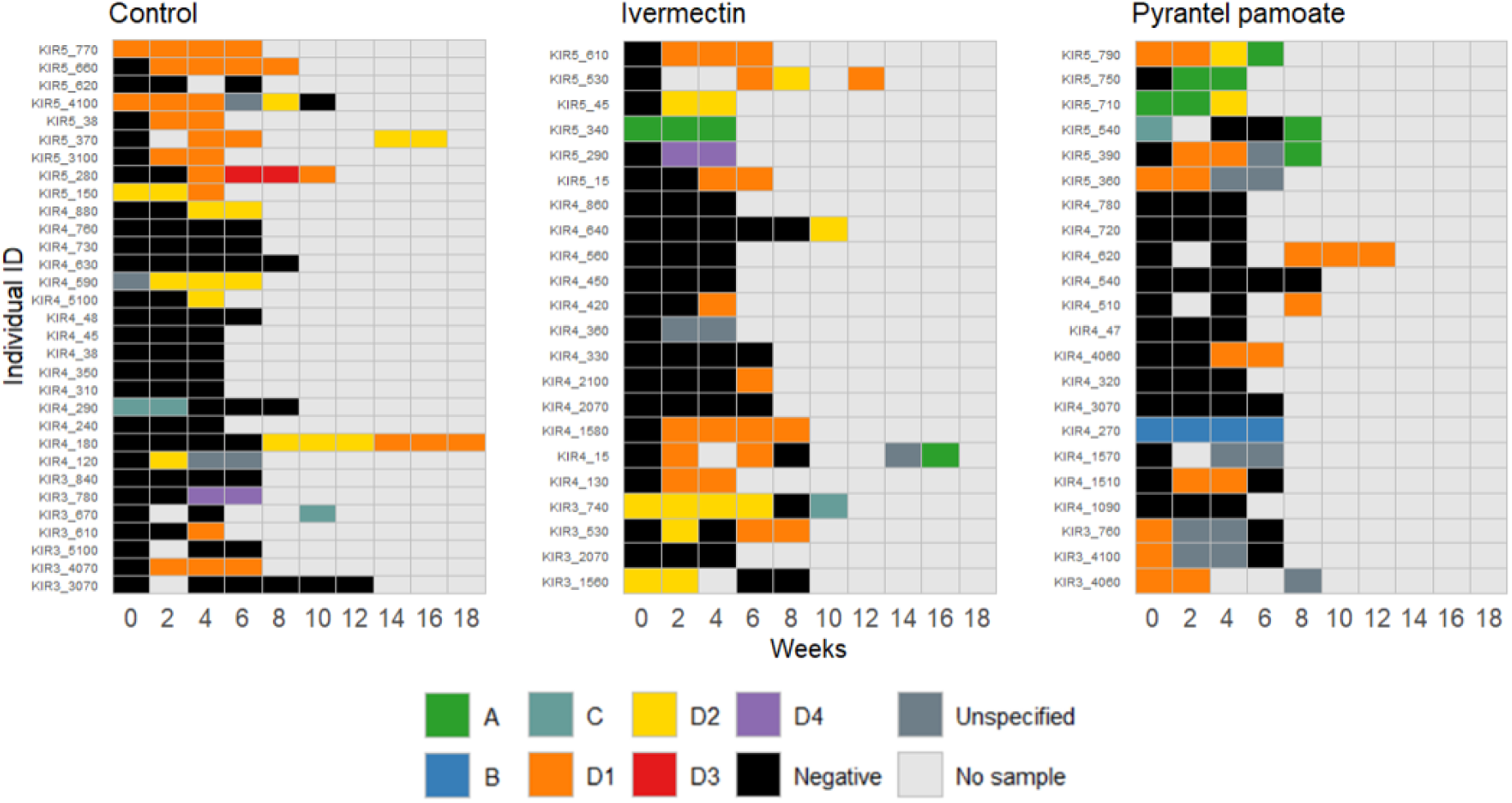
Visualization of individual *Bartonella* infection patterns across 10 sampling time points for each treatment group (control, ivermectin and pyrantel pamoate) in wild *Mastomys natalensis* populations. Rasters display the infection status (positive with respective genogroup, negative or no sample) for each individual at each time point.

## 4. Discussion

Co-infections, particularly between helminths and microparasites, play a crucial role in pathogen transmission dynamics, with potential implications for public health. However, experimental evidence from natural populations, including perturbation experiments, remains scarce, especially on the African continent. To address this gap, we conducted a laboratory experiment with wild-caught rodents to assess the efficacy of two anthelmintic treatments (ivermectin and pyrantel pamoate) followed by a field experiment in Tanzania to investigate how helminth infections influence microparasite dynamics in *Mastomys natalensis*. This rodent species is a common pest in sub-Saharan Africa and an important host of various pathogens, some of which can be transmitted to humans. Our results showed that both treatments significantly reduced gastrointestinal nematode infections, with pyrantel pamoate being more effective than ivermectin. Interestingly, pyrantel pamoate treatment was associated with a decline in Morogoro virus (MORV) seropositivity over time, suggesting a potential link between helminth infection and host susceptibility to viral infection. In contrast, *Bartonella* infection probability remained unaffected by anthelmintic treatment, indicating that helminths play a limited role in shaping *Bartonella* dynamics.

Our study identified six different helminth genera and two families, including five nematodes (*Trichuris mastomysi*, *Syphacia* sp., *Gongylonema* sp., *Protospirura* sp. and Trichostrongylidae) and three cestodes (*Hymenolepis* spp., Taenia sp. and Davaineidae). Among these, *T. mastomysi* and Davaineidae were the most prevalent, with prevalence rates of 63.2% and 57.9% respectively. These helminths are commonly found in rodents across Africa (Brouat et al. 2007, Diagne et al. 2016) and the high diversity of gastrointestinal parasites observed in this study is consistent with previous research on *M. natalensis* in Tanzania (Vanden Broecke et al. 2021). However, the prevalence rates we observed differed from those reported by Vanden Broecke et al. (2021), likely due to factors specific to each site, such as environmental conditions, which can affect parasite prevalence and abundance. Similarly, Brouat et al. (2007) highlighted that local environmental factors are important in determining helminth prevalence and diversity. Alternatively, variations in sample size and the effects of anthelmintic treatment in our study may have influenced the detectability of certain helminths, potentially leading to lower observed prevalence rates. Notably, this study presents the first report of *Taenia* cysts in *M. natalensis*. Although this is not unexpected, given that *Taenia* spp. are known to use small rodents as intermediate hosts (Thaikoed et al. 2024), their presence raises concern due to the zoonotic potential of certain *Taenia* species. This finding highlights the need for further research on the role of *M. natalensis* in the transmission cycle of *Taenia* species.

Both anthelmintic treatments, ivermectin and pyrantel pamoate, effectively reduced the prevalence of gastrointestinal nematodes, although their efficacy varied. Pyrantel pamoate was significantly more effective, especially in reducing *T. mastomysi* infections, while ivermectin showed only partial efficacy. This result is surprising, as pyrantel pamoate is generally not considered highly effective against *Trichuris* spp. (Wilson, 2012). One reason for the lower efficacy of ivermectin could be the relatively low dose used in this study to minimize potential side effects. However, since treatment did not significantly affect body weight, indicating no adverse effects on animal health or well-being, future studies could consider using a higher dose of ivermectin to potentially improve efficacy (Wahid et al. 1998). Neither treatment significantly reduced cestode prevalence, which was expected, as both drugs primarily target nematodes (Wahid et al. 1998).

Our models showed that treatment with pyrantel pamoate was associated with a decrease in the likelihood of being positive for Morogoro virus-specific antibodies (MORVab) over time. This suggests that reducing helminth infections with pyrantel pamoate may decrease the likelihood of maintaining MORV infection, indicating that helminths might affect host susceptibility to the virus. This finding contradicts our expectations, as MORVab are generally thought to persist for life based on laboratory experiments (Borremans et al. 2015). However, findings from laboratory studies do not always match what we see in natural populations (Mariën et al. 2017). On the other hand, ivermectin had no significant effect on seropositivity, likely due to its limited efficacy against *T. mastomysi*. A stronger reduction in helminths might be needed to observe an effect on microparasite dynamics (Fenton et al. 2013). Overall, the differences between the groups were small, and the large confidence intervals suggest caution in interpretation. Future studies with larger sample sizes, direct testing for MORV RNA, and a deeper investigation into how helminths affect viruses could provide conclusive answers.

Unlike its effects on MORV, anthelmintic treatment did not affect the likelihood of *Bartonella* infection. This suggests that helminths do not strongly influence host susceptibility to *Bartonella*, despite previous observational studies indicating a potential positive association (Vanden Broecke et al. 2023). However, we found that heavier animals were more likely to be infected. Since body weight correlates with age in *M. natalensis* (Leirs et al. 1990), older animals may have had more time to be exposed to *Bartonella* (Vanden Broecke et al. 2023). Similarly, an age effect was also observed for helminth infections in *M. natalensis* (Vanden Broecke et al. 2021), which could explain the previously reported association between helminths and *Bartonella* infection. Increased risk of infection in heavier animals may also be linked to body size, as larger animals can carry more ectoparasites and have a greater surface area, increasing their exposure while moving through their environment. Indeed, body size has been shown to influence ectoparasite abundance in other rodent species, such as *Oligoryzomys nigripes* (Fernanda et al. 2015). In addition, we found that infection probability varied between grids, likely due to environmental factors, like vegetation type and density, which can influence ectoparasite presence and abundance (Santos et al. 2018, Smith et al. 2023).

Notably, this spatial variation was observed not only in infection probability but also in strain distribution, with genogroup A almost exclusively found in KIR5. This could be due to differences in vector species, which may influence the local adaptation of *Bartonella* strains (Chomel et al. 2009). In fact, the majority of the *Bartonella* strains identified in this study belonged to *Bartonella elizabethae*, a rodent-associated species linked to human diseases such as endocarditis (Kosoy et al. 2012, Hayman et al. 2013), further underscoring *M. natalensis* as an important reservoir for zoonotic pathogens. Additionally, genogroup C may represent a novel *Bartonella* species, as it did not cluster with previously identified isolates, though it showed some similarity *B. tribocorum*. However, further studies are needed to confirm this classification and to fully characterize this group (Kosoy et al. 2018).

In conclusion, our findings highlight the complex interactions between helminths and microparasites in a natural setting. While helminth reduction influenced Morogoro virus seropositivity, no significant effect was observed for *Bartonella*, suggesting that microparasite responses to helminth infection may be species-specific. Perturbation experiments in wild populations present inherent challenges, including the need for highly effective antiparasitic treatments and the difficulty of recapturing individuals over time, which can reduce sample sizes and introduce uncertainties in the results. Further research with larger sample sizes, direct detection of microparasites, and a deeper understanding of the underlying mechanisms is necessary to fully understand helminth-microparasite interactions.

## Ethics statement

All experimental procedures were approved by the Ethics Committee for Animal Experimentation of the University of Antwerp (2021-04) and adhered to the EEC Council Directive 2010/63/EU. The research also adhered to the animal ethics guidelines from the Sokoine University of Agriculture’s ‘Code of Conduct for Research Ethics’ (Revised version of 2012) and the guidelines by Sikes and Gannon (2016).

## Availability of data

The sequences generated in this study will be submitted to the GenBank repository upon acceptance of the manuscript.

## CRediT authorship contribution statement

**Marre van de Ven**: Conceptualization, Investigation, Data Curation, Formal Analysis, Writing – Original draft, Writing – Review & Editing. **Bram Vanden Broecke**: Formal Analysis, Writing – Review & Editing. **Alexis Ribas**: Investigation **Christopher Sabuni**: Resources. **Herwig Leirs**: Funding acquisition. **Joachim Mariën**: Conceptualization, Supervision, Methodology, Project Administration, Writing – Review & Editing.

## Acknowledgments

We thank the staff at the Institute of Pest Management (Sokoine University of Agriculture, Morogoro, Tanzania), particularly Geoffrey Sabuni, Shabani Lutea, Omary Kibwana, Ramadhani Iddy and Baraka Edson, for their invaluable assistance during the fieldwork and dissections.

## Supplementary

**Table S1:** Results from generalized linear models (GLM) analysing the effects of anthelmintic treatment (Ivermectin, Pyrantel pamoate vs control), sex, age and their interaction on nematode infection in *Mastomys natalensis*. Significant effects (p < 0.05) are underlined.

| *Nematodes ~ treatment \* sex + treatment \* age* |  |  |  |
| --- | --- | --- | --- |
| **Coefficients** | **Estimate ± SE** | **z-ratio** | **p-value** |
| (intercept) | 1.566 ± 1.618 | 0.968 | 0.333 |
| Ivermectin vs control | -1.552 ± 2.329 | -0.666 | 0.505 |
| Pyrantel pamoate vs control | -4.786 ± 2.257 | -2.121 | <u>0.034</u> |
| Sex – Male | 17.670 ± 2168.000 | 0.008 | 0.994 |
| Age | -0.006 ± 0.013 | -0.503 | 0.615 |
| Ivermectin : Male | -20.820 ± 2168.000 | -0.010 | 0.992 |
| Pyrantel pamoate : Male | -18.360 ± 2168.000 | -0.008 | 0.993 |
| Ivermectin : Age | 0.022 ± 0.023 | 0.951 | 0.342 |
| Pyrantel pamoate : Age | 0.022 ± 0.019 | 1.193 | 0.233 |
| *Nematodes ~ treatment + sex + age* |  |  |  |
| (intercept) | 1.600 ± 1.127 | 1.420 | 0.156 |
| Ivermectin vs control | -1.821 ± 0.787 | -2.316 | <u>0.021</u> |
| Pyrantel pamoate vs control | -3.548 ± 0.925 | -3.838 | <u><0.001</u> |
| Sex – Male | -0.726 ± 0.681 | -1.066 | 0.286 |
| Age | 0.004 ± 0.009 | 0.489 | 0.625 |
| *Nematodes ~ treatment* |  |  |  |
| (intercept) | 1.674 ± 0.629 | 2.661 | 0.008 |
| Ivermectin vs control | -1.875 ± 0.773 | -2.425 | <u>0.015</u> |
| Pyrantel pamoate vs control | -3.409 ± 0.888 | -3.840 | <u><0.001</u> |

**Table S2:**
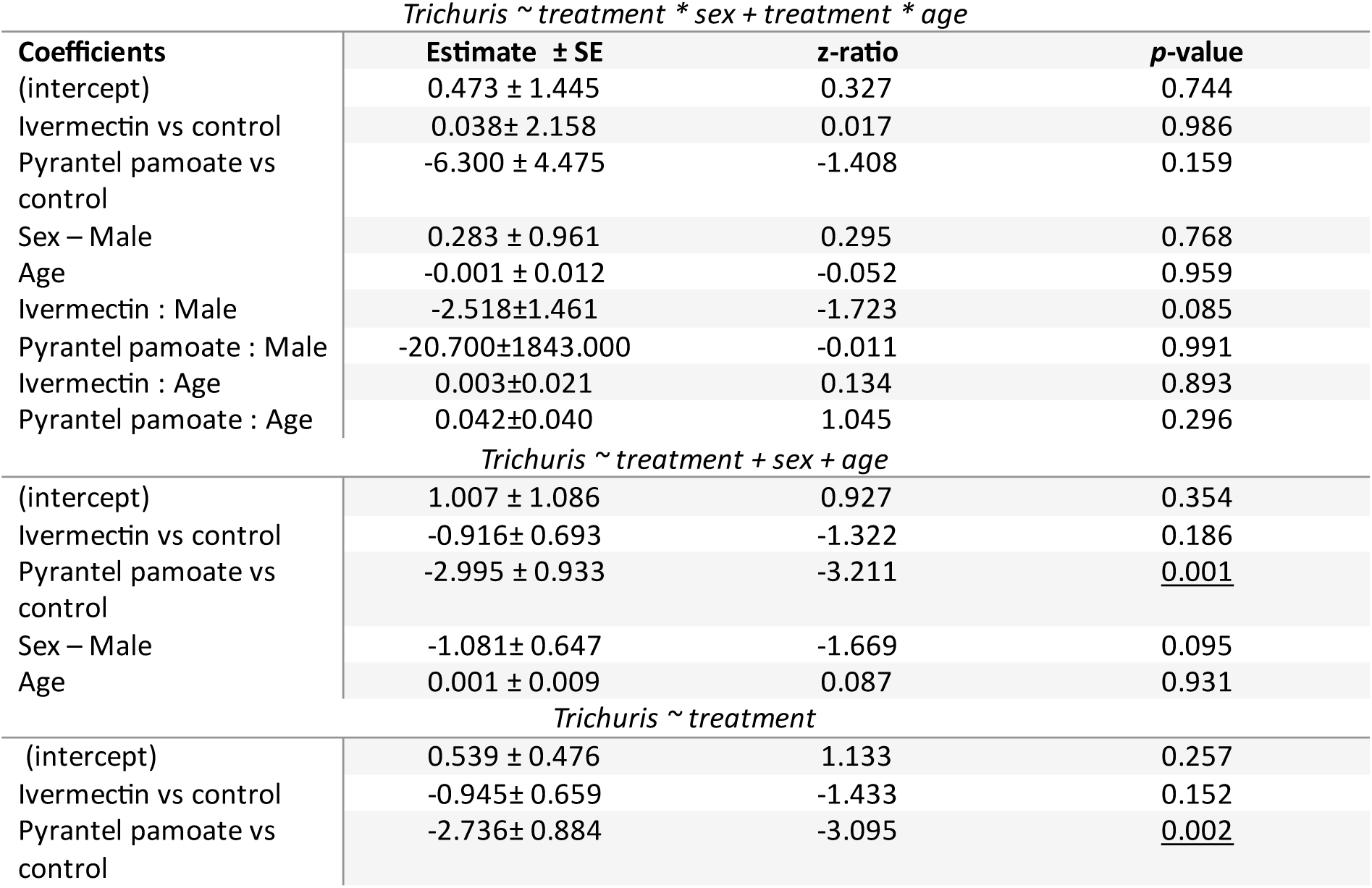
Results from generalized linear models (GLM) analysing the effects of anthelmintic treatment (Ivermectin, Pyrantel pamoate vs control), sex, age and their interaction on *Trichuris mastomysi* infection in *Mastomys natalensis*. Significant effects (p < 0.05) are underlined.

**Table S3:**
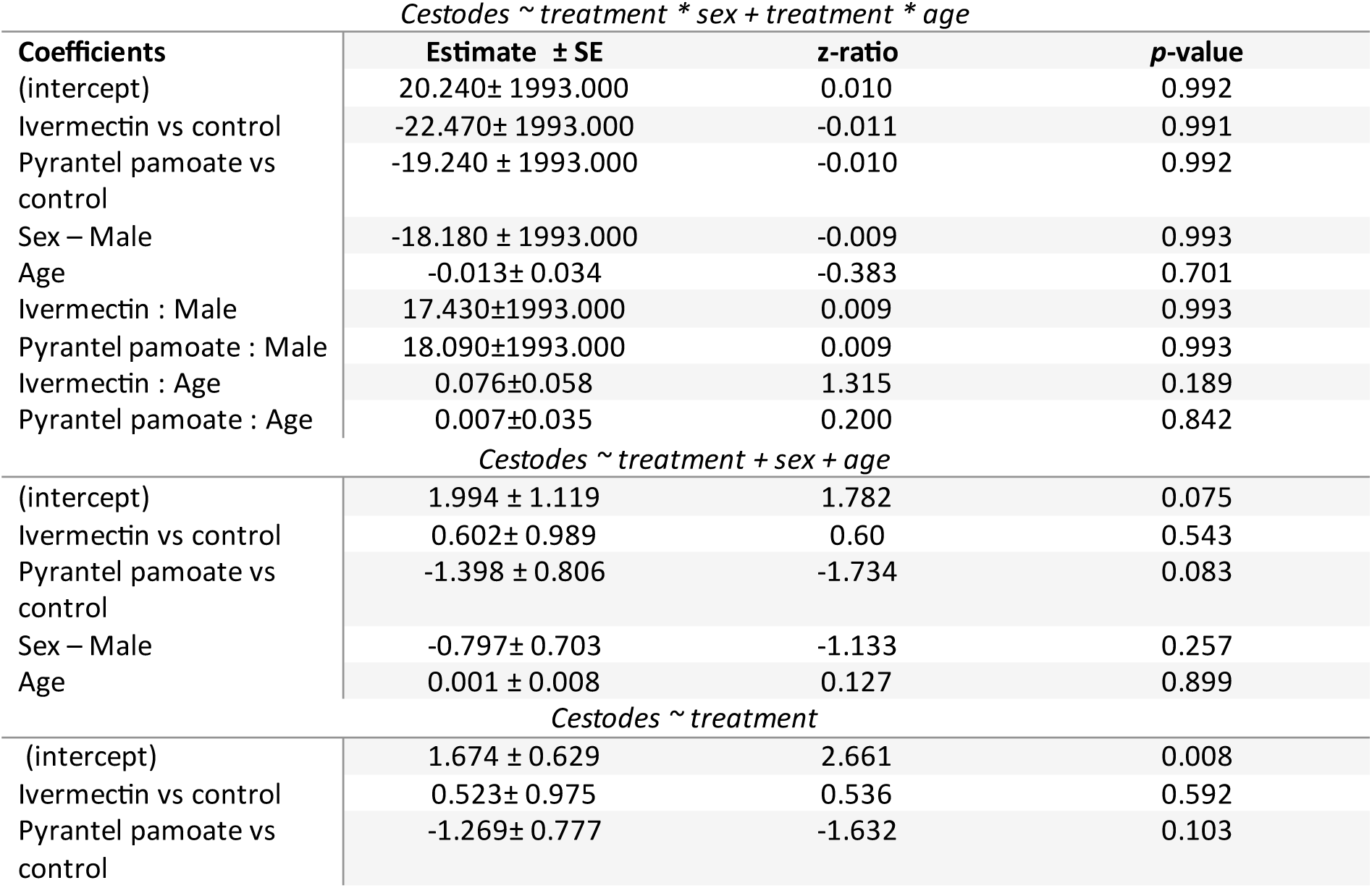
Results from generalized linear models (GLM) analysing the effects of anthelmintic treatment (Ivermectin, Pyrantel pamoate vs control), sex, age and their interaction cestode infection in *Mastomys natalensis*. Significant effects (p < 0.05) are underlined.

| *Cestodes ~ treatment \* sex + treatment \* age* |  |  |  |
| --- | --- | --- | --- |
| **Coefficients** | **Estimate ± SE** | **z-ratio** | **p-value** |
| (intercept) | 20.240± 1993.000 | 0.010 | 0.992 |
| Ivermectin vs control | -22.470± 1993.000 | -0.011 | 0.991 |
| Pyrantel pamoate vs control | -19.240 ± 1993.000 | -0.010 | 0.992 |
| Sex – Male | -18.180 ± 1993.000 | -0.009 | 0.993 |
| Age | -0.013± 0.034 | -0.383 | 0.701 |
| Ivermectin : Male | 17.430±1993.000 | 0.009 | 0.993 |
| Pyrantel pamoate : Male | 18.090±1993.000 | 0.009 | 0.993 |
| Ivermectin : Age | 0.076±0.058 | 1.315 | 0.189 |
| Pyrantel pamoate : Age | 0.007±0.035 | 0.200 | 0.842 |
| *Cestodes ~ treatment + sex + age* |  |  |  |
| (intercept) | 1.994 ± 1.119 | 1.782 | 0.075 |
| Ivermectin vs control | 0.602± 0.989 | 0.60 | 0.543 |
| Pyrantel pamoate vs control | -1.398 ± 0.806 | -1.734 | 0.083 |
| Sex – Male | -0.797± 0.703 | -1.133 | 0.257 |
| Age | 0.001 ± 0.008 | 0.127 | 0.899 |
| *Cestodes ~ treatment* |  |  |  |
| (intercept) | 1.674 ± 0.629 | 2.661 | 0.008 |
| Ivermectin vs control | 0.523± 0.975 | 0.536 | 0.592 |
| Pyrantel pamoate vs control | -1.269± 0.777 | -1.632 | 0.103 |

**Table S4:** Results from a generalized linear mixed model (GLMM) analysing the effects of anthelmintic treatment (Ivermectin, Pyrantel pamoate vs control), time, their interaction, sex and bodyweight on the presence of Morogoro-virus specific antibodies in wild *Mastomys natalensis* populations. Significant predictors (p < 0.05) are underlined.

| *Morogoro virus ~ treatment \* time + sex + body weight + grid (1 \| ID)* |  |  |  |
| --- | --- | --- | --- |
| **Coefficients** | **Estimate ± SE** | **z-ratio** | **p-value** |
| (intercept) | -7.309±2.289 | -3.194 | 0.001 |
| Ivermectin vs control | -0.556±1.457 | -0.381 | 0.703 |
| Pyrantel pamoate vs control | 0.827±1.402 | 0.590 | 0.555 |
| Time | 0.408±0.240 | 1.705 | 0.088 |
| Sex – Male | 0.419± 1.021 | 0.411 | 0.681 |
| Body weight | 0.111± 0.052 | 2.133 | <u>0.033</u> |
| Grid – KIR4 | 1.024± 1.383 | 0.740 | 0.460 |
| Grid – KIR5 | 1.042± 1.495 | 0.697 | 0.486 |
| Ivermectin : Time | -0.062± 0.355 | -0.174 | 0.862 |
| Pyrantel pamoate : Time | -0.829 ± 0.384 | -2.162 | <u>0.031</u> |

**Table S5:**
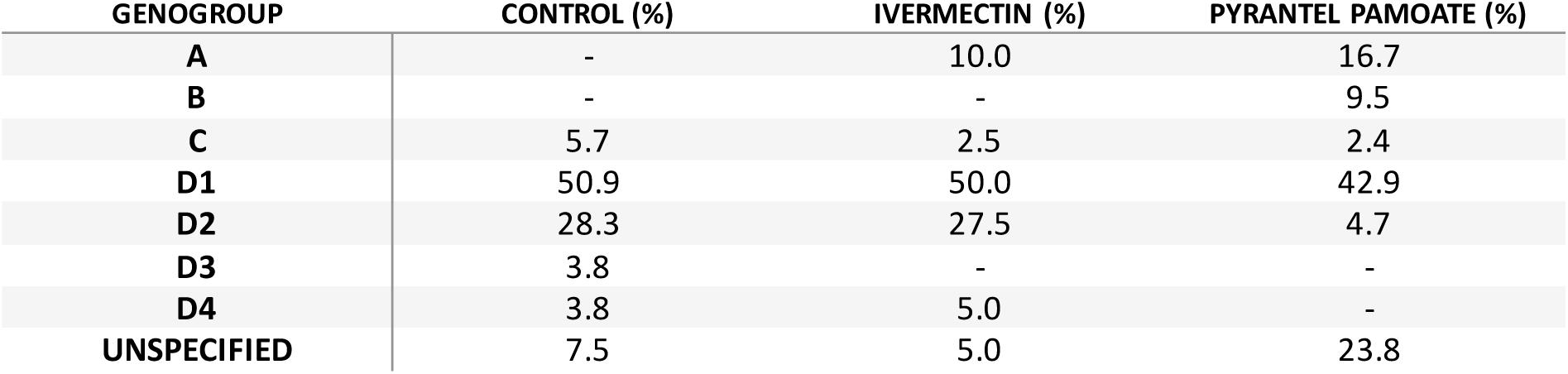
Proportions of samples belonging to each *Bartonella* genogroup across anthelmintic treatment groups (control, ivermectin and pyrantel pamoate). A dash (“-”) indicates that the genogroup was not observed in the respective treatment group.

| GENOGROUP | CONTROL (%) | IVERMECTIN (%) | PYRANTEL PAMOATE (%) |
| --- | --- | --- | --- |
| A | - | 10.0 | 16.7 |
| B | - | - | 9.5 |
| C | 5.7 | 2.5 | 2.4 |
| D1 | 50.9 | 50.0 | 42.9 |
| D2 | 28.3 | 27.5 | 4.7 |
| D3 | 3.8 | - | - |
| D4 | 3.8 | 5.0 | - |
| UNSPECIFIED | 7.5 | 5.0 | 23.8 |

**Table S6:** Results from generalized linear mixed models (GLMM) analysing the effects of anthelmintic treatment (ivermectin and pyrantel pamoate), time, their interaction, sex and bodyweight on *Bartonella* infection in wild *Mastomys natalensis*. Significant predictors (p < 0.05) are underlined.

| *Bartonella* ~ treatment * time + sex + body weight + grid + (1\|ID) |  |  |  |
| --- | --- | --- | --- |
| Coefficients | Estimate ± SE | z-ratio | p-value |
| (intercept) | -8.552 ± 2.343 | -3.650 | <u><0.001</u> |
| Ivermectin vs control | -0.364 ± 1.331 | -0.274 | 0.784 |
| Pyrantel pamoate vs control | 3.154 ± 1.398 | 2.255 | <u>0.024</u> |
| Time | 1.038 ± 0.308 | 3.365 | <u><0.001</u> |
| Sex – Male | -0.118 ± 0.787 | -0.150 | 0.880 |
| Body weight | 0.225 ± 0.063 | 3.570 | <u><0.001</u> |
| Grid – KIR4 | -4.234 ± 1.430 | -2.962 | <u>0.003</u> |
| Grid – KIR5 | 1.875 ± 1.163 | 1.612 | 0.107 |
| Ivermectin x Time | 0.323 ± 0.408 | 0.792 | 0.428 |
| Pyrantel x Time | -0.501 ± 0.384 | -1.302 | 0.193 |
| *Bartonella* ~ treatment + time + sex + body weight + grid + (1\|ID) |  |  |  |
| (intercept) | -8.984 ± 2.656 | -3.382 | <u><0.001</u> |
| Ivermectin vs control | 0.551 ± 1.031 | 0.534 | 0.593 |
| Pyrantel pamoate vs control | 2.280 ± 1.183 | 1.927 | 0.054 |
| Time | 1.065 ± 0.281 | 3.787 | <u><0.001</u> |
| Sex – Male | -0.052 ± 0.873 | -0.059 | 0.953 |
| Body weight | 0.233 ± 0.070 | 3.331 | <u><0.001</u> |
| Grid – KIR4 | -4.732 ± 1.720 | -2.752 | <u>0.006</u> |
| Grid – KIR5 | 2.188 ± 1.335 | 1.639 | 0.101 |

**Figure S1:**
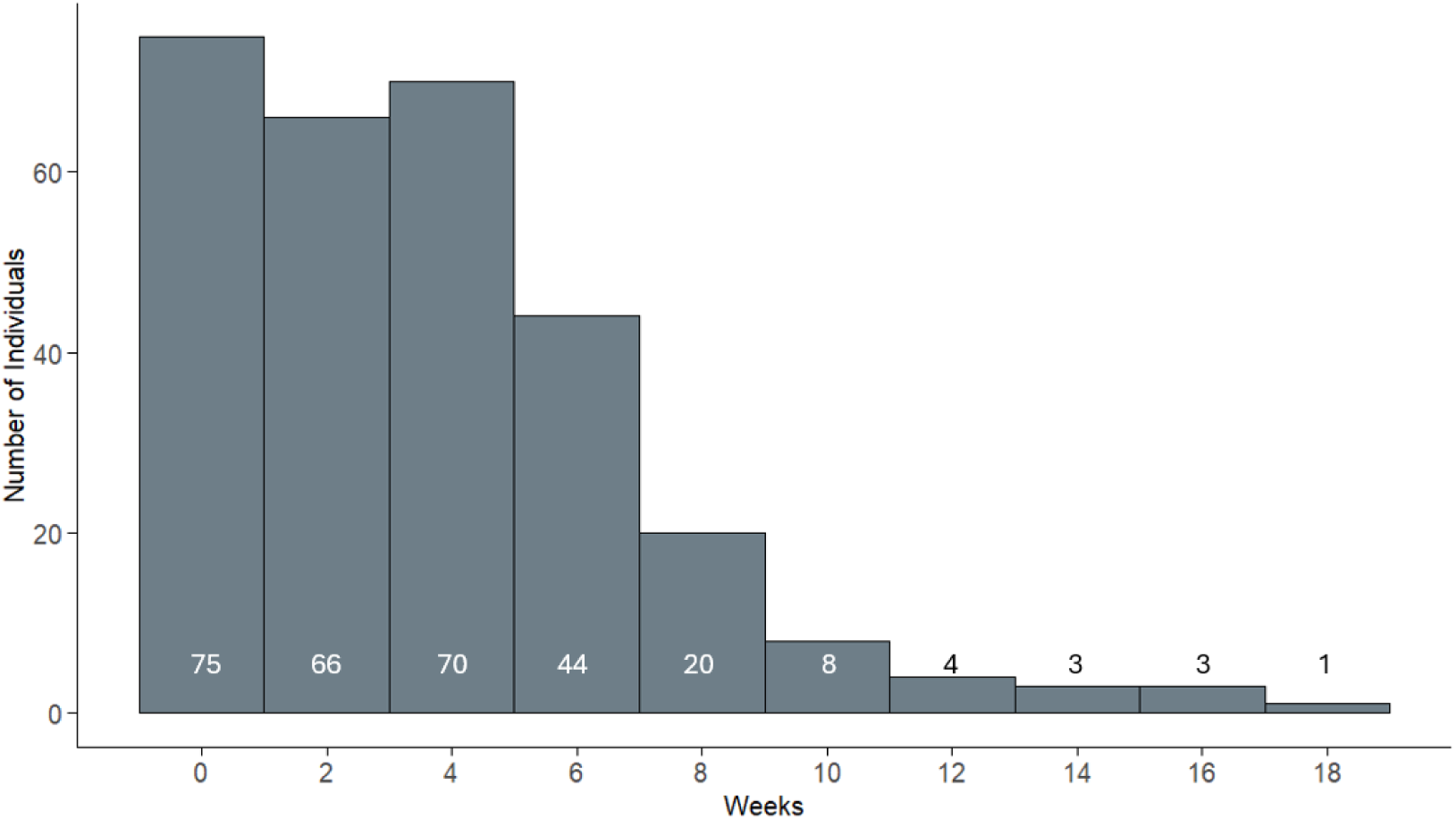
Histogram showing the capture frequency of individual *Mastomys natalensis* (n=75) included in the capture-mark-recapture analysis over an 18-week period. The first capture was set as time 0, with subsequent time points calculated relative to this initial capture (in weeks). The exact number of individuals captured at each time point is indicated at the bottom of the bars.

**Figure S2:**
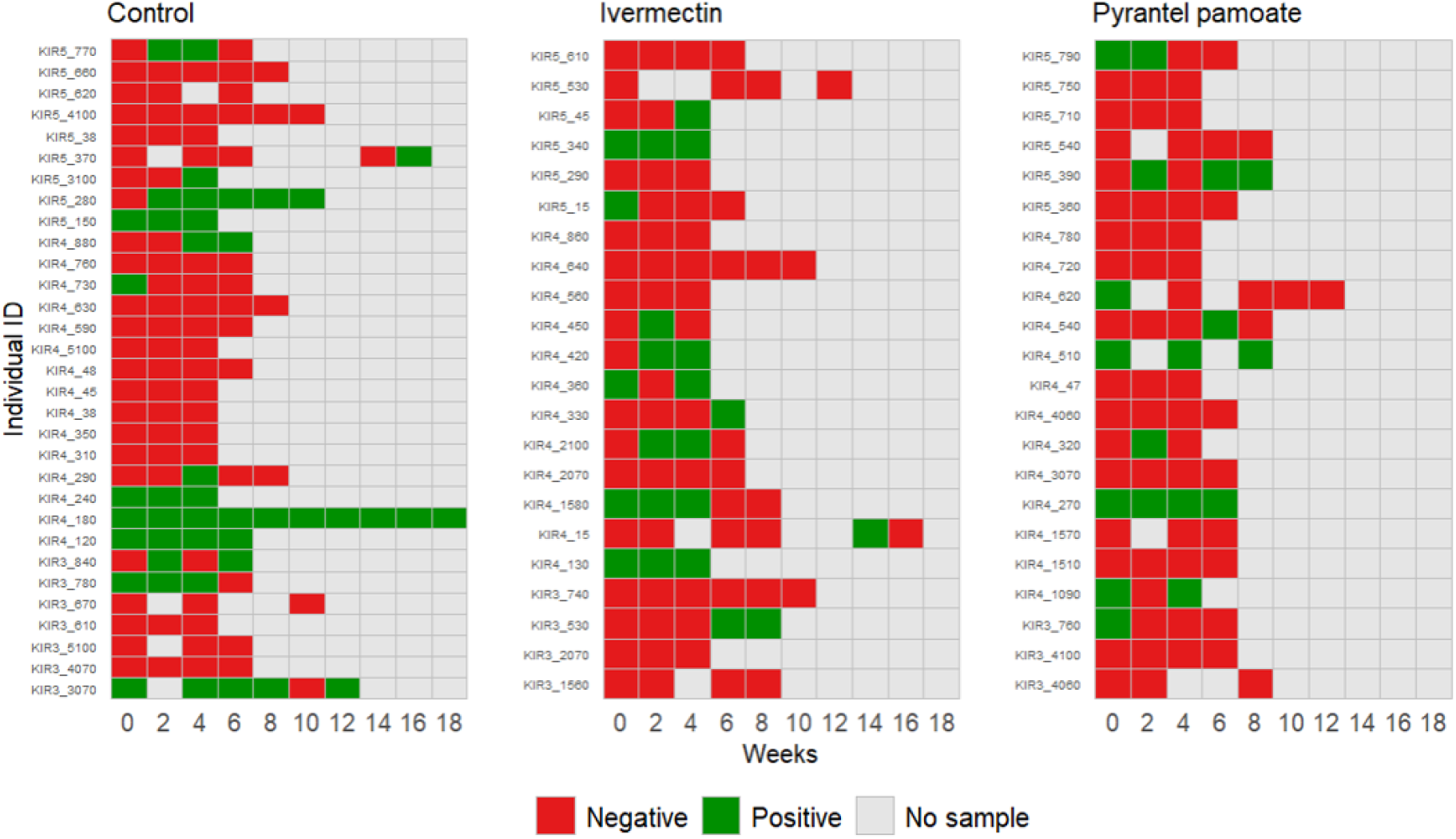
Visualization of individual Morogoro virus antibody patterns across 10 sampling time points for each treatment group (control, Ivermectin and Pyrantel pamoate) in wild *Mastomys natalensis* populations. Rasters display the antibody status (positive, negative or no sample) for each individual at each time point.

